# A Cone–Plate Bioreactor for Applying Physiologically Derived Shear Stress Waveforms to Study Endothelial Mechanotransduction and Cardiac Cell Signaling

**DOI:** 10.64898/2026.05.31.729120

**Authors:** Matthew C. Watson, Erica C. Kemmerling, Lauren D Black

## Abstract

Fluid shear stress is a critical regulator of endothelial cell function and cardiovascular development, yet *in vitro* platforms often lack the ability to reproduce physiologically relevant, time-dependent flow environments with quantitative precision. Here, we present the design and validation of a macro-scale cone–plate bioreactor engineered to deliver controlled steady and pulsatile shear stress waveforms to endothelial monolayers and engineered tissues. The system integrates a geometry optimized to minimize secondary flow effects, a feedback-controlled motor capable of reproducing complex waveforms, and a viscosity-informed control framework to account for shear-dependent fluid behavior.

Using this platform, endothelial cells were exposed to steady and physiologically derived pulsatile shear stresses. Cells exhibited increased alignment and eccentricity under shear, confirming biologically relevant mechanical stimulation. While pulsatile shear did not significantly alter endothelial neuregulin-1 expression, exogenous administration studies revealed a nonlinear, dose-dependent increase in cardiomyocyte proliferation. Furthermore, co-culture experiments demonstrated that shear-conditioned endothelial cells promote cardiomyocyte proliferation, suggesting a mechanotransduction-mediated paracrine signaling mechanism. Together, these results establish a versatile and quantitatively controlled platform for studying cardiovascular mechanobiology. This device enables systematic investigation of shear-dependent cellular responses and provides a foundation for integrating co-culture systems and three-dimensional engineered tissues under physiologically relevant hemodynamic conditions.

## INTRODUCTION

The endothelium forms a dynamic interface between circulating blood and cardiovascular tissues, where it plays a central role in regulating transport, mechanotransduction, and tissue homeostasis. Endothelial cell phenotype and function are strongly influenced by fluid shear stress, a mechanical cue arising from blood flow that varies in magnitude, directionality, and waveform throughout the cardiac cycle ^1-3^. Both steady and pulsatile shear stresses modulate endothelial alignment, cytoskeletal organization, gene expression, and signaling pathways, including nitric oxide production and mechanoresponsive transcriptional programs ^1, 3-8^. These effects are particularly important during cardiac development, where flow-induced forces contribute to morphogenesis and congenital disease progression.

A key challenge in cardiovascular mechanobiology is the controlled, quantitative reproduction of physiologic shear stress conditions in experimental systems. While in vivo approaches provide physiologically relevant flow environments, they cannot decouple hemodynamic forces from systemic biological complexity or precisely control local mechanical parameters ^9-11^. Conversely, in vitro platforms enable controlled interrogation of mechanobiological responses but often lack the ability to reproduce the time-dependent, spatially uniform shear stress waveforms present in the developing and adult cardiovascular system. Existing in vitro flow platforms span microfluidic devices and macroscale bioreactors. Microfluidic systems offer precise control over geometry and biochemical gradients but are typically limited to low Reynolds number regimes, restricting their ability to reproduce physiologically relevant pulsatile and inertial flow conditions ^1^. Macroscale systems—including parallel-plate and cone–plate devices— provide improved access to physiologic shear stress magnitudes and temporal dynamics. Parallel-plate systems generate well-defined laminar shear but are generally limited to unidirectional flow profiles ^5, 12^. Cone–plate devices enable relatively uniform shear stress across a large area and are well-suited for implementing time-varying and oscillatory shear waveforms, although they may introduce secondary flow effects near geometric boundaries ^12, 13^.

Despite these advances, current systems are limited in their ability to simultaneously provide (i) physiologically accurate waveform control, (ii) spatially uniform shear fields over biologically relevant areas, and (iii) compatibility with co-culture and engineered tissue platforms. Furthermore, few studies explicitly integrate non-Newtonian fluid behavior into device operation, despite its importance for accurately translating physiological shear stresses to in vitro conditions ^13-16^.

Here, we present the design and validation of a macro-scale cone–plate bioreactor engineered to deliver customizable, physiologically derived shear stress waveforms to endothelial monolayers and engineered tissues. The system incorporates (i) a geometry optimized to minimize secondary flows over defined culture regions, (ii) a feedback-controlled motor system capable of reproducing complex pulsatile waveforms, and (iii) a viscosity-informed control framework that accounts for shear-dependent fluid properties.

Using this platform, we evaluate endothelial responses to steady and pulsatile shear stress and investigate mechanically mediated endothelial–cardiomyocyte signaling, focusing on neuregulin-1 (Nrg1) as a candidate paracrine regulator of cardiomyocyte proliferation ^17-21^. This work establishes a device-centric framework for studying cardiovascular mechanobiology and provides a foundation for extending controlled shear stimulation to engineered cardiac tissues.

## MATERIALS AND METHODS

### Cone-plate shearing bioreactor

The design of our cone-plate bioreactor is a large scale version of cone-plate shearing viscometers (Figure 1 A-B). There were several design constraints. First, the cone-plate geometry needed to be large enough such that flow over cell regions would be approximately unidirectional, and cells needed to be placed where secondary flows were negligible. Secondary flows and eddy formulations are prevalent near the far walls and the cone vertex in cone-plate shearing geometries. Our design featured a 0.5 degree cone, truncated at the vertex to minimize grinding with the base-plate, and a 13 inch diameter base-plate. A minimum gap height of 0.5mm between the truncated cone and plate was set by a 0.5 inch diameter cylindrical plug. Six 22mm x 22mm x 0.2mm cut-outs for the installation of cultured cells were machined into the baseplate along a10 cm bolt-circle. Here, the direction of flow varied by less than 4% over the cells and cut-out locations were far enough away from the cone truncation and the walls that secondary flow effects are negligible (Figure 1 C).

**Figure 1.**
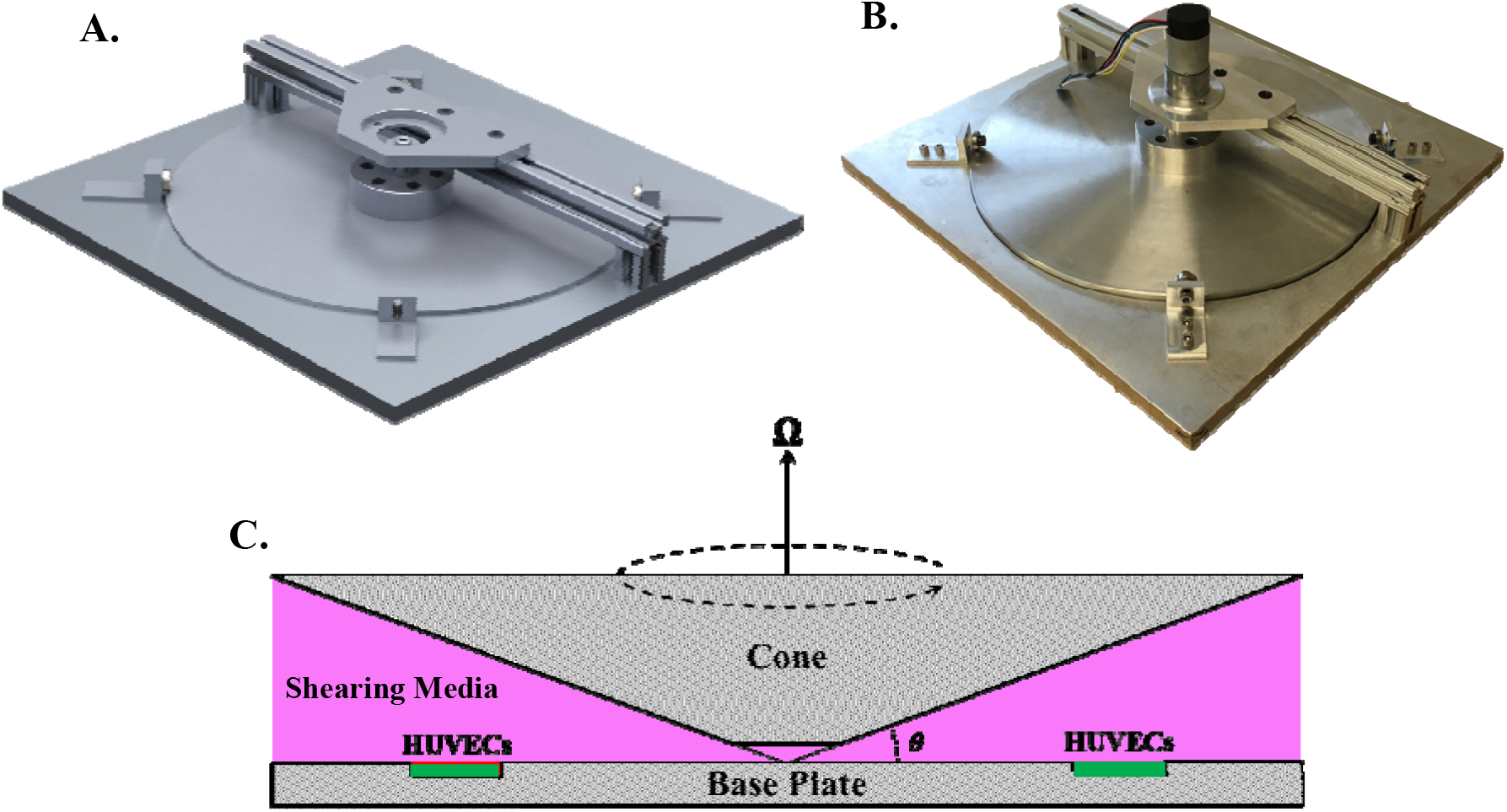
(A) Computer aided design isometric rendering of our cone-plate shearing bioreactor and (B) an isometric image of the manufactured and assembled bioreactor. (C) Exaggerated schematic showing a cross section of the operating device. Cell locations 10 cm away from the cone vertex, the truncated cone has a minimum gap height of 0.5 mm, and a cone angle (θ) of 0.5º. The angular velocity is denoted by Ω.

Our bioreactor assembly consisted of multiple components and a drive unit, and it was necessary that all components in contact with cells be aseptic and biocompatible. Additionally, oxidation was a concern, as our bioreactor operated in an incubated environment maintained at 37ºC, 5% CO_2_, and 95% humidity. To prevent oxidation and maintain biocompatibility, the base-plate and cone were aluminum, the gap-height plug was polytetrafluoroethylene (PTFE), and all fasteners were stainless steel. All components in contact with cells and the incubator were steam autoclaved at 121 ºC for sterilization prior to experiments.

### Drive unit

The bioreactor drive unit consisted of a DC gearmotor and encoder (Pololu) controlled by an Arduino. A custom designed proportional integral derivative (PID) controller drove the motor and provided both steady shear stress waveforms and complex patient specific shear stress waveforms. Coefficients were tuned prior to the start of an experiment (code available upon request). The motor shaft was coupled directly to the cone, while the controller remained outside of the incubator. Sensitive electronics within the motor housing prevented the steam autoclave from being a viable sanitation method. Instead, the motor and wires were wiped with 70% ethanol before operation within the incubator.

### Shearing medium

The device was filled with basal endothelial cell medium (EBM-2, Lonza) supplemented with essential growth factors (EGM-2) to shear endothelial cells (Figure 1 C). To realistically model wall shear stresses experienced by endothelial cells in a blood-filled environment, it was necessary to know the viscous properties of both blood and the shearing medium. We utilized data collected by Brooks *et al*. to account for the shear thinning behavior of blood, and we performed rheometry to determine the shear medium viscosity as a function of increasing shear rate. The TA Instrument’s AR-G2 rheometer was used to measure the viscosity of our shearing medium at shear rates from 10 – 800 s^-1^. A steel cone geometry with an angle of 2º was used. The rheometer and the medium were kept at 37ºC throughout the viscosity measurement acquisition. A solvent trap was used to prevent evaporation. Deionized water is a Newtonian fluid, and to remove operating error, viscosity measurements of deionized water were used to normalize the viscosity of the cell medium. A shear-thinning behavior was observed, with viscosity decreasing with increasing shear rates and leveling off at shear rates larger than 500 s^-1^. A piecewise polynomial was used to fit the viscosity data (Figure 2 A).

**Figure 2:**
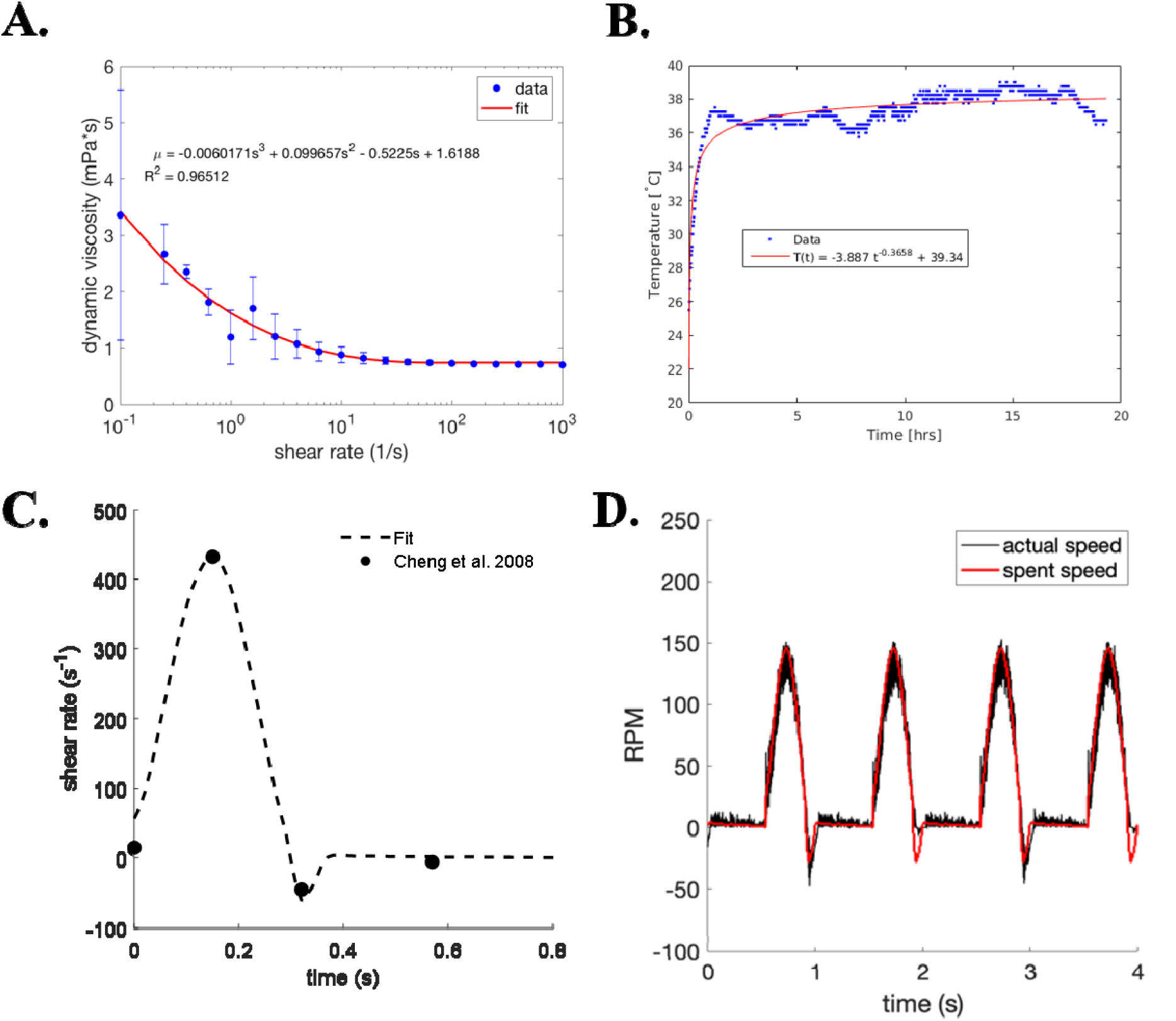
Viscosity and Motor Data. **(A)** Viscosity measurements and fit for endothelial cell media. **(B)** Motor temperature data and fit of bioreactor operating at max speed for 18 hours. **(C)** Shear rate data and fit from data of healthy flow through an adult aorta from Chen et al. **(D)** Sent speeds (red) and measured output speeds (black) of our bioreactor.

### Shear stress waveforms

Shear stress waveforms were extracted from CFD models of patient specific healthy aorta models by Chen *et al*. (Figure 2 C) ^22^. Briefly, time dependent velocity profiles were extracted using the open source software package, WebPlotDigitizer. From the velocity profiles, a linear fit of the velocities near the walls was used to determine wall shear rates (equation 1).

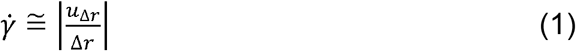

Here, 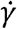 is the shear rate, Δ*r* is the distance from the wall, and *u*_Δ*r*_ is the velocity at Δ*r*. Flow rate waveforms were also extracted using WebPlotDigitizer. A cubic spline was used to fit each flow rate waveform, and the respective flow-rate waveform shapes were used to fit the wall shear rates (Figure 2 C). The viscosity of blood at varying shear rates was determined by a fit of data collected by Brooks *et al*. for 40% hematocrit blood ^23, 24^, and the dependence of shear stress on time of blood (|*τ*_*blood*_ |) were calculated by multiplying the shear rate 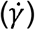 and the viscosity fit of blood 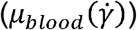 (equation 2).

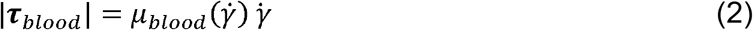

The wall shear stress of blood calculated from the aortic flow data was used to calculate the shear rate waveform to drive the motor. Since both blood viscosity and the shearing medium viscosity were dependent on shear rate, an iterative method was used to calculate the motor speed. Briefly, an initial guess for the shearing medium viscosity (*µ*_0_) was used to calculate the first iteration of shear rate for the motor 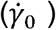 (equation 3). Then the viscosity estimate (*µ*_*shear*_) was adjusted and a new value for the shear rate 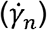 was calculated (equation 4). A maximum of 100 iterations were performed to converge viscosity and shear rate and set the motor’s speed.

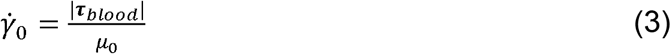

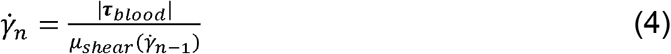

The healthy wall shear stress waveform featured a maximum shear stress of 13.4 dynes/cm^2^ and a minimum of −3.1 dynes/cm^2^ at a frequency of 1.25 Hz. In experiments, endothelial cells were exposed to either the healthy waveforms or a steady shear stress of 3.1 dynes/cm^2^, which was the time-averaged shear stress of the healthy waveform.

### Endothelial cell culture for shearing experiments

Human umbilical vein endothelial cells (HUVECs) that had been lentivirally infected with CytoLight Green (Essen BioScience, Cat No 4453; Ann Arbor, MI) were used for all experiments. HUVECs were kept in flasks and on tissue culture plastic plates that had been coated with a gelatin fibronectin coating for at least 2 hours in an incubator or at room temperature overnight, in a sterile environment. The gelatin fibronectin coating was made by mixing 0.02% gelatin (Sigma-Aldrich; St. Louis, MO) in sterile water with 1mg/ml fibronectin (Thermofisher; Waltham, MA). After plating, HUVECs were kept in sterile incubators at 37°C with 5% CO_2_ and cultured in optimized growth media, consisting of EBM-2 (Lonza; Walkersville, MD) and EGM-2 singleQuot kit supplemental and growth factors (Lonza; Walkersville, MD). Media was changed every 2-3 days and HUVEC passage numbers 4-8 were used for experiments. Prior to the start of shearing experiments, HUVECs were passaged and seeded at a density of 25,000 cells/cm^2^ onto 22mm x 22mm glass coverslips coated with gelatin fibronectin. After seeding, HUVECs were cultured until confluency (1-2 days) before coverslips were installed into the shearing bioreactor. HUVECs plated on gelatin fibronectin coated coverslips were kept as a control.

### Cardiomyocyte culture for exogenous NRG1 and Shearing experiments

Neonatal rat cardiomyocytes were isolated from Sprague-Dawley rat pups at p1. Neonatal rats were placed on ice and euthanized by decapitation. For each pup, the chest wall was carefully opened, and the heart was removed. Then, hearts were minced, and tissues underwent 7 x 7 min digestions in collagenase type II (Worthington Biochemical Corp, Lakewood, NJ) and sterile PBS supplemented with 20 mM glucose. Cells were counted with a hemocytometer and seeded onto tissue culture plates at a density of 50,000 cells/cm^2^. The culture medium contained 15% fetal bovine serum (FBS) in Dulbecco’s Modified Eagle Medium (DMEM) with 1% penicillin-streptomycin and was changed every 2 days.

### Immunohistochemistry

Cellular monolayers were fixed by immersion in 4% paraformaldehyde (158127, Sigma-Aldrich) at 4°C for 10 minutes. Samples then underwent three washes in 1X phosphate buffered saline (PBS), before being permeabilized using 0.05% solution of triton X in 1X PBS for ten minutes at room temperature. Samples then underwent three washes in 1X PBS. A 5% donkey serum in 1% bovine serum albumin solution (BSA) (A-9418, Sigma-Aldrich) was used to block samples at room temperature for at least one hour. Cells were stained with primary antibodies for either α-actinin (Sigma; A7811), neuregulin-1 (Abcam; ab125708), or Ki67 (Abcam; ab15580) at concentrations of 1:200 in 1% BSA overnight at 4°C. Following three washes in 1X PBS, a 1:400 concentration of secondary antibody (Thermo Fisher) in 1% BSA was applied to samples for 1 hour at room temperature. All cell staining also included nuclear staining using 4′,6′-diamidino-2-phenylindole (DAPI) (Thermo Fisher Scientific; D-1306). DAPI was added during the secondary antibody staining. Samples were again washed three times before imaging.

Custom pipelines in CellProfiler (Broad Institute) were used to observe cell morphology and the measure the percentages of cells expressing α-actinin, neuregulin-1, or Ki67. For all image quantification, at least five images per sample were used. Morphological pipelines counted all the DAPI positive nuclei and objects expressing green fluorescent protein (GFP). Then, the pipeline created a mask of DAPI positive nuclei. The mask was then applied to GFP expressing objects, creating a new image resulting in only GFP objects containing a nucleus on which morphological properties were measured. Similar pipelines assessed the percentages of cells expressing GFP, α-actinin, neuregulin-1, or Ki67. Briefly, all DAPI positive nuclei and objects expressing GFP or α-actinin were counted. A mask was created from either GFP or α-actinin objects and applied to the DAPI nuclear stain images. This subsequently counted all nuclei contained within GFP or α-actinin positive cells. Similarly, the pipeline counted all positive stains for either neuregulin-1 or Ki67. A mask created from nuclei expressing either GFP or α-actinin was then applied to neuregulin-1 or Ki67 objects. Subsequently, this counted all nuclei expressing both GFP and neuregulin-1 or α-actinin and Ki67.

### Exogenous neuregulin on cardiomyocytes

We exposed neonatal rat cardiomyocytes cultured in a twelve well tissue culture plastic plate to exogenous neuregulin-1 to assess the proliferative effects of neuregulin-1. Freshly isolated neonatal rat cardiomyocytes were seeded at a density of 50,000 cells/cm^2^. Each well was cultured with 2 mL of serum-free media containing either 200 ng/mL of neuregulin-1, 100 ng/mL of neuregulin-1, or no neuregulin-1. After 72 hours, cells were fixed with 4% paraformaldehyde, labeled with anti-cardiac α-actin and anti-Ki67 (a proliferative marker), and imaged as previously described. Cells without neuregulin administration were also fixed. Custom pipelines developed in CellProfiler (Broad Institute, Cambridge, MA) assessed proliferation of cells as described in above.

### Effects of conditioned medium from sheared endothelial cells on primary cardiomyocytes

To assess whether ECs under pulsatile shear impacted CM phenotype, we collected culture medium from ECs. HUVECs were seeded on coverslips as described above and allowed to reach confluency before being cultured in the cone-plate system for 48 hours under pulsatile shear. Conditioned medium was collected and mixed in a 1:1 ratio with normal culture medium for neonatal rat cardiomyocytes (see above for formulation). This medium was then added to wells containing CMs seeded at 50,000 cells/cm^2^ on tissue culture plastic. 1:1 conditioned medium from ECs cultured under static conditions was used as a control. After 24 or 72 hours of culture in EC-conditioned media, the Cardiomyocytes were fixed in 4% paraformaldehyde, stained with anti-cardiac α-actin and anti-Ki67, and imaged as previously described. Custom pipelines developed in CellProfiler (Broad Institute, Cambridge, MA) assessed proliferation of cells as described above.

### Statistics

A minimum of 3 samples were used for all experimental analysis. Replicate values are indicated in the figure legends. Two tailed student t-tests or a one-way ANOVA with Tukey post hoc testing were performed using GraphPad Prism v8 for comparisons and are indicated in the figure legends. Significance was determined by p values ≤ 0.05.

## RESULTS

### Validation of cone-plate shearing bioreactor

We first needed to evaluate the performance of our motor. To determine maximum motor temperature, we filled the bioreactor with the shearing medium and ran it at maximum speed within the incubator for 18 hours. A thermocouple in contact with the motor casing output temperature readings every minute. After 18 hours the device maintained operation, and the motor temperature leveled off to 39ºC (Figure 2 B). Next, we needed to validate that the motor output matched the input waveform. The device was filled with the shear medium and speed readings were sampled at 1000 Hz validating that our device could output the desired pulsatile waveform (Figure 2 D).

To validate that our bioreactor was capable of culturing endothelial cells under shear stress, we applied a steady shear stress of 3.1 dynes/cm^2^ for 24 hours. Endothelial cells expressing GFP remained attached to coverslips for the duration of the shearing cycle and aligned in the direction of flow (Figure 3 A-B). Cell eccentricity and spread area were calculated using a custom pipeline in CellProfiler (Broad Institute). Eccentricity was significantly increased in endothelial cells cultured under shear compared to static controls (p < 0.05) (Figure 3C), but we observed no significant difference in cell spread area (Figure 3D).

**Figure 3:**
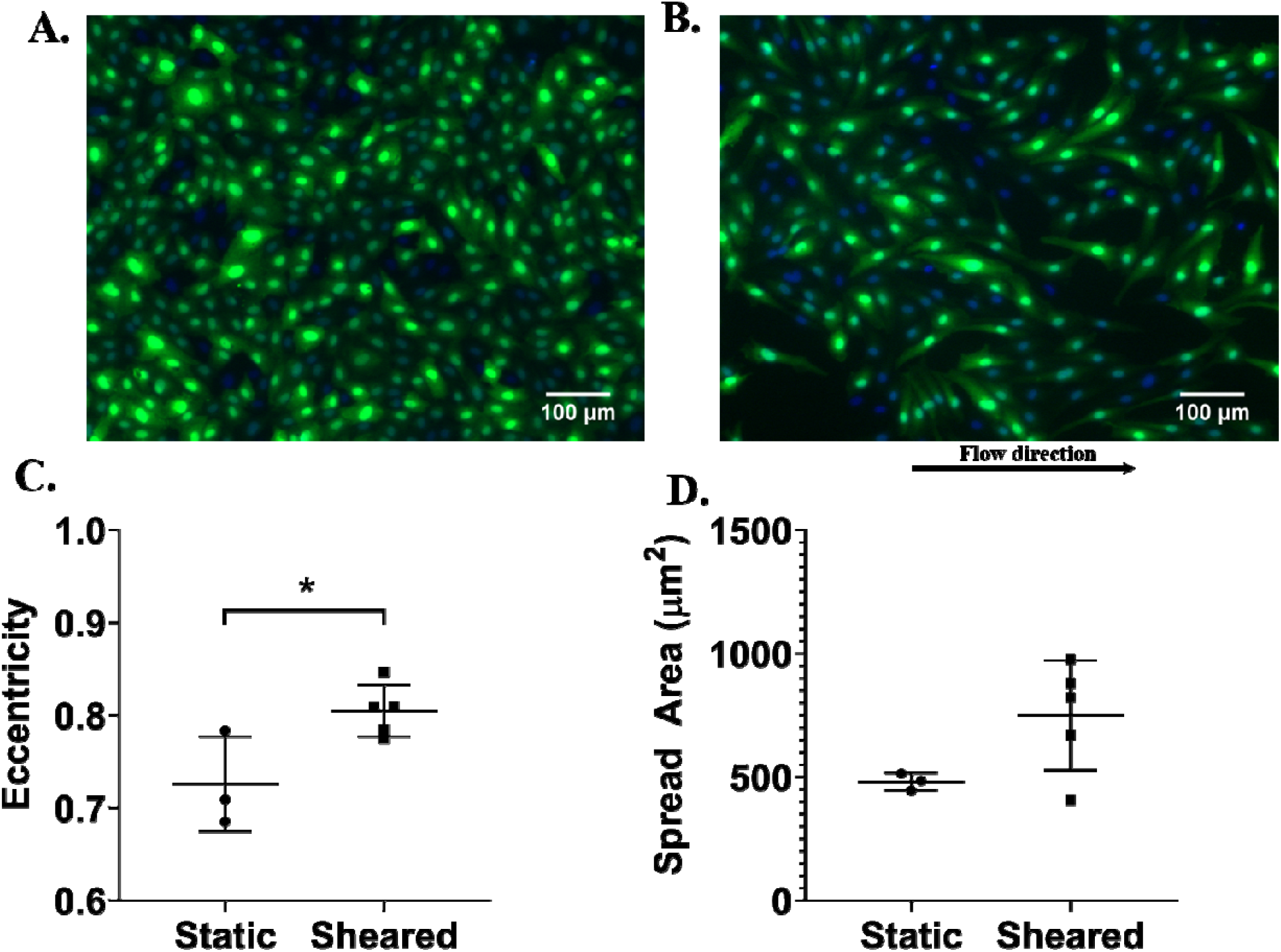
Steady Shear Culture. Representative images of GFP (green) expressing endothelial cells and nuclei (blue) under static conditions **(A)** or steady shear of 3.1 dynes/cm^2^ **(B)**. Cell morphologies were analyzed using a custom script in CellProfiler. A significant increase cell eccentricity was observed in endothelial cells cultured under shear stress **(C)**; however, no significant change in cell spread area was observed **(D)**. (n ≥ 3; * = p ≤ 0.05; comparisons by two-tailed student t-test; scale bar = 100 µm)

### Endothelial cells cultured under pulsatile shear stress

Next, we applied pulsatile shear stress mimicking healthy aortic flow to endothelial cells. Endothelial cells were cultured under pulsatile shear stress for 24 hours. After 24 hours, cells were fixed and stained for neuregulin-1. Cell attachment was an issue for pulsatile experiments (p < 0.01) (Figure 4 A-B), as culture under pulsatile shear stress resulted in a significant decrease in cell density compared to static culture (Figure 4 C). Endothelial cells exposed to pulsatile shear stress were significantly more eccentric than the static control (p < 0.05) (Figure 4 D). No significant change was observed for positive expression of neuregulin-1 in endothelial cells cultured under pulsatile shear stress compared to the static control (Figure 4 E).

**Figure 4:**
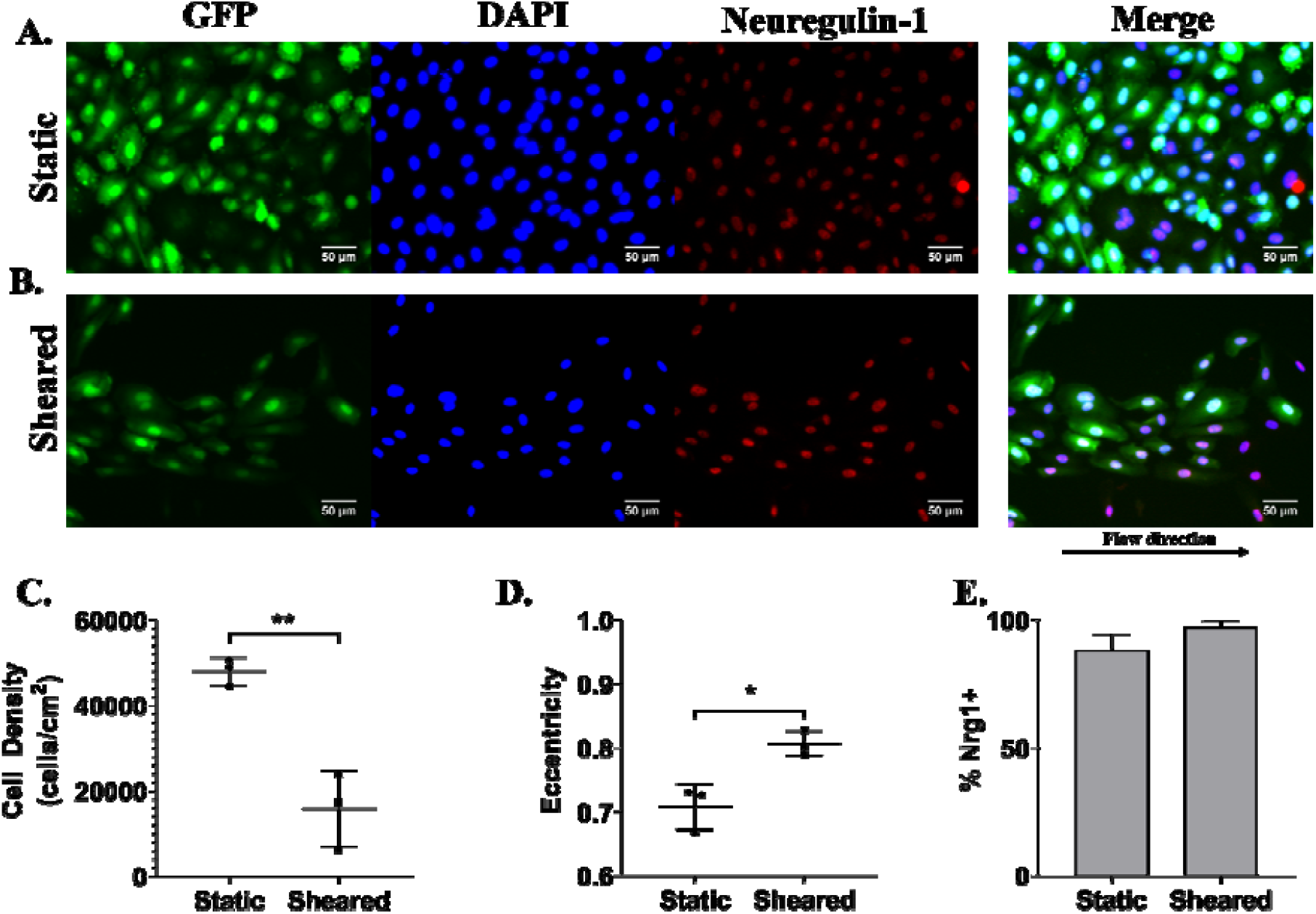
Pulsatile Shear Culture. Representative images of GFP (green) and neuregulin-1 (red) expressing endothelial cells and nuclei (blue) under static conditions **(A)** or pulsatile shear stress (**B**). Cell densities, morphologies, and protein expression were analyzed using a custom script in CellProfiler. The cell density significantly decreased after culture under pulsatile shear stress **(C)**. Cell cultured under pulsatile shear stress had higher eccentricities **(D)**. No significant difference was observed in the percentage of cells expressing neuregulin-1 (E). (n = 3; p = * ≤ 0.05; comparisons by two-tailed student t-test; scale bar = 50 µm)

### Exogenous neuregulin-1 promotes cardiomyocyte proliferation in vitro

We were interested in examining how different concentrations of exogenous neuregulin-1 affect cardiomyocyte proliferation. Freshly isolated cardiomyocytes were cultured with increasing concentrations of neuregulin-1 and stained for cardiac α-actin (cardiomyocyte marker) and Ki-67 (proliferative marker) after 72 hours (Figure 5 A-C). A significant increase in the percentage of cells expressing α-actinin was observed in cells cultured with a neuregulin-1 concentration of 100 ng/mL compared to cells cultured without neuregulin-1 (p < 0.05) (Figure 5 D); however, no significant difference was observed in the fold change of α-actinin expressing cardiomyocytes at 72 hours from α-actinin expressing cardiomyocytes without exogenous neuregulin-1 treatment at 0 hours (Figure 5 E). Staining for Ki-67 revealed a significant increase in the percentage of proliferating cardiomyocytes treated with 100 ng/mL of neuregulin-1 compared to both cardiomyocytes cultured without neuregulin-1 and cardiomyocytes cultured with 200 ng/mL of neuregulin-1 (p < 0.001) (Figure 5 F). Additionally, a significant increase was observed in the fold change of expression of Ki-76 at 72 hours from cardiomyocyte expression without exogenous neuregulin-1 at the start of the experiment (p < 0.01) (Figure 5 G).

**Figure 5:**
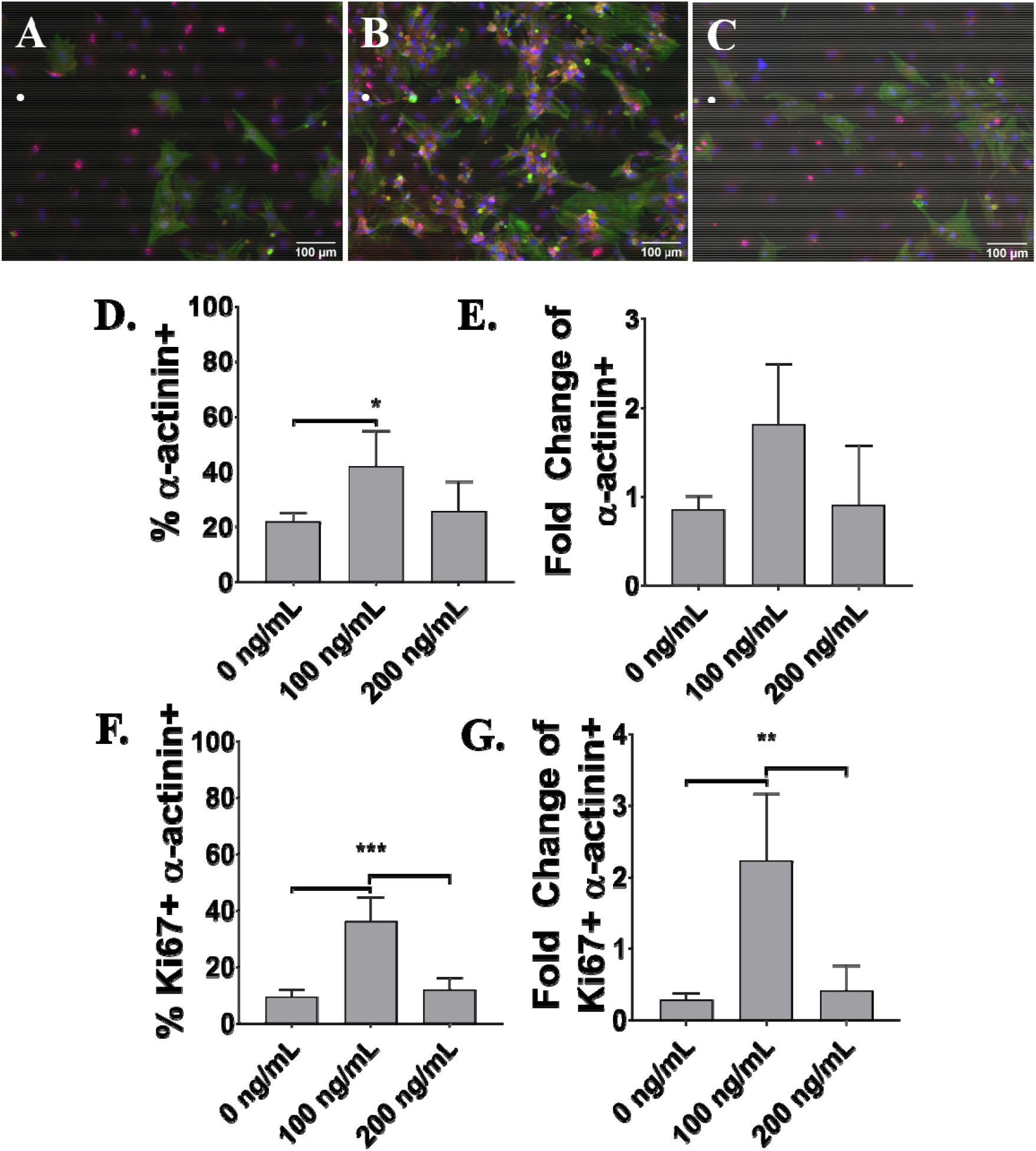
Exogenous Neuregulin-1. Representative images Ki67 (red) and α-actinin (green) expressing cells (blue) cultured without neuregulin-1 **(A)** or with neuregulin-1 at concentrations of 100 ng/mL **(B**), or with 200 ng/mL **(C)** at 72 hours. A significant increase in the percentage of cells expressing α-actinin after 72 hours **(D)**. No significance was observed in the fold change of α-actinin positive cells at 72 hours compared to cells without neuregulin administration at 0 hours **(E)**. Both the percentage of Ki67 expressing cardiomyocytes **(F)** and the fold change of Ki67+ cardiomyocytes at 72 hours compared to cells without neuregulin administration at 0 hours were significantly higher than cells cultured without neuregulin and cells cultured with 200 ng/mL.(**G**). (n = 4; * = p < 0.05, ** = p < 0.01, *** = p < 0.001; comparisons by one-way ANOVA with Tukey post hoc test; scale bar = 100 µm).

### Pulsatile shear stress on endothelial cells promotes cardiomyocyte proliferation

Next, we investigated the effects endothelial cells cultured under shear stress have on the proliferative capability of cardiomyocytes. In order to separate direct effects of shearing on the cardiomyocytes themselves, we chose to culture only the endothelial cells either statically or under pulsatile shear for 48 hours and then create conditioned media in a 1:1 ratio with standard cardiomyocyte medium. After 24 hours and 72 hours of culture in EC-conditioned medium from static and pulsatile sheared conditions, cardiomyocytes were stained for both α-actinin and Ki67 (Figure 6 A-D). No significant differences were observed in the percentage of α-actinin expression between sheared and static conditioned media at 24 hours or 72 hours (Figure 6 E); however, at the 72 hour timepoint, there was a significant increase in the percentage cells expressing both α-actinin and Ki67 for sheared conditioned medium compared to static conditioned medium (p < 0.05) (Figure 6 F). This suggests that culture of endothelial cells under pulsatile shear stress conditions promotes cardiomyocyte proliferation through paracrine signaling, but more work is necessary to elucidate the biological mechanisms involved.

**Figure 6:**
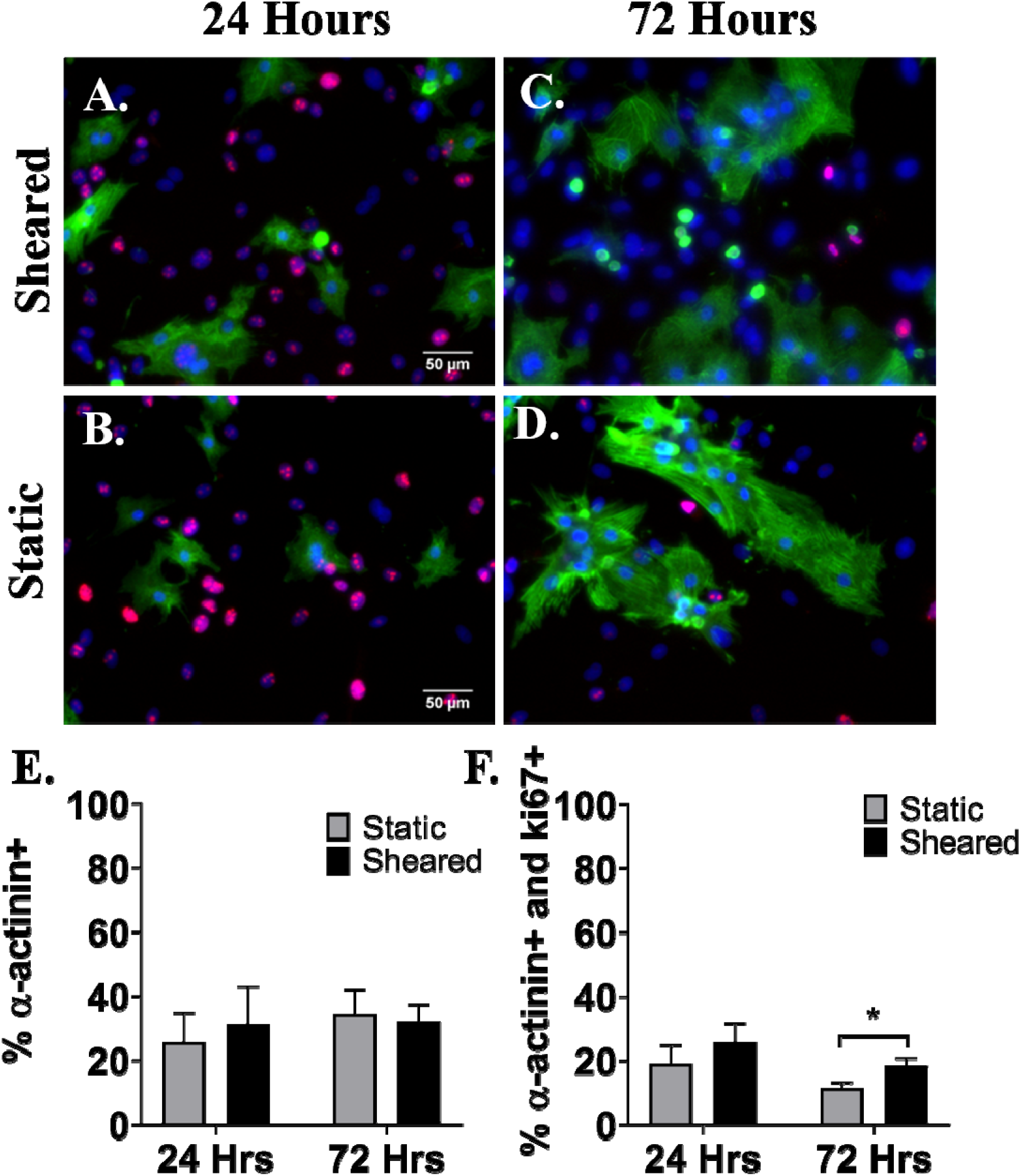
Sheared ECs paracrine effects on CMs. Representative images of neonatal rat cells stained for nuclei (blue), α-actinin (green), and Ki67 (red) at 24 hours after application of conditioned medium derived from ECs under pulsatile shear stress **(A)** or static EC culture **(B)**, or at 72 hours after application of conditioned medium derived from ECs under pulsatile shear stress **(C)** and or static EC culture **(D)**. Protein expression was analyzed using custom scripts in CellProfiler. Percentages of α-actinin positive cells after 24 hours and 72 hours after conditioned medium addition **(E)**. Percentages of α-actinin and Ki67 positive cells after 24 hours and 72 hours of conditioned medium exposure **(F)**. (n = 3; * = p < 0.05; comparisons by multiple student t-tests with Holm-Sidak post hoc testing; scale bar = 50 µm)

## Discussion

In this study, we developed and validated a cone–plate bioreactor designed to apply controlled, physiologically relevant shear stress waveforms to cultured cells, addressing key limitations in existing *in vitro* flow platforms. The device integrates geometric optimization, closed-loop motor control, and viscosity-aware waveform translation to enable reproducible delivery of both steady and pulsatile shear stresses. Importantly, this approach enables dynamic, waveform-resolved control of the mechanical microenvironment, which is critical for faithfully modeling cardiovascular physiology ^1, 6^.

A central engineering consideration in cone–plate systems is the balance between achieving spatially uniform shear stress and minimizing secondary flow artifacts. Prior work has demonstrated that cone–plate geometries can introduce non-ideal flow features, particularly near the cone apex and outer boundaries ^12, 13^. By selecting a shallow cone angle (0.5°), truncating the cone apex, and positioning culture regions away from these boundary effects, our design minimizes these contributions while preserving a large usable culture area. The observed alignment and increased eccentricity of endothelial cells under shear (Figures 3 and 4) confirm that the device generates biologically relevant shear stresses consistent with classical endothelial responses to flow ^4, 5^, supporting its validity as a mechanobiological platform.

Importantly, our system enables implementation of physiologically derived, time-dependent shear stress waveforms, extending beyond the steady or sinusoidal conditions commonly used in *in vitro* studies. The ability to reproduce waveform features such as flow reversal and transient peaks is particularly relevant for modeling cardiovascular environments, including the embryonic heart, where shear stress is inherently pulsatile ^2, 9, 25^. While the applied waveforms were derived from adult aortic flow, their magnitudes fall within the range reported for embryonic systems ^9, 25^, suggesting that the device can be adapted to developmental models with appropriate waveform inputs. These capabilities reinforce the importance of capturing temporal complexity in shear stress, as endothelial signaling pathways are highly sensitive to waveform characteristics ^1, 8^.

A notable feature of the present approach is the explicit incorporation of shear-dependent fluid viscosity into waveform generation. Blood exhibits shear-thinning behavior, and neglecting this property can lead to discrepancies between intended and delivered shear stresses, particularly under pulsatile conditions ^23, 26^. By iteratively solving for motor speed based on both target shear stress and the rheological properties of the culture medium ^23, 26^, our system improves fidelity in mapping physiological conditions to *in vitro* experiments. This represents an important design consideration that is often omitted in comparable platforms.

From a biological perspective, previous investigations *in vivo* suggest that neuregulin-1 promotes cardiomyocyte proliferation, but that elevated or ectopic levels can drive pathological outcomes such as hyper-trabeculation ^19, 21, 27^. In the present study, endothelial cells cultured under pulsatile shear did not exhibit significant changes in Nrg1 expression relative to static controls, despite clear morphological responses. However, it is important to note that Nrg1 expression was assessed at a single time point, and the absence of an observed difference may reflect the temporal dynamics of Nrg1 regulation rather than a true lack of shear responsiveness. Mechanosensitive signaling pathways in endothelial cells often exhibit transient, time-dependent activation profiles, with rapid induction of transcriptional programs such as KLF2 mediated through flow-sensitive mechanotransduction pathways ^8, 28, 29^. Thus, it is possible that changes in Nrg1 expression occurred outside of the selected assessment window. In contrast, exogenous Nrg1 stimulation produced a nonlinear proliferative response in cardiomyocytes, with maximal effects observed at 100 ng/mL and attenuation at higher concentrations, consistent with prior reports of dose-dependent and context-specific Nrg1 signaling ^19, 21, 27^.

Importantly, the conditioned media experiments (Figure 6) demonstrate that endothelial cells exposed to pulsatile shear stress can promote cardiomyocyte proliferation through soluble signaling factors. In these studies, cardiomyocytes were not directly exposed to shear; rather, they were treated with media conditioned by endothelial cells under pulsatile flow. This result strengthens the conclusion that endothelial mechanotransduction is sufficient to modulate cardiomyocyte behavior via paracrine signaling, independent of direct mechanical stimulation.

The apparent discrepancy between unchanged Nrg1 expression and increased cardiomyocyte proliferation in response to conditioned media may reflect several factors. First, as noted above, Nrg1 expression may be temporally regulated and not captured at the measurement time point. Second, endothelial cells secrete a broad array of mechanosensitive paracrine factors—including nitric oxide, endothelin-1, prostacyclin, and angiocrine growth factors—that collectively regulate cardiomyocyte growth, survival, and function through endothelial–cardiomyocyte crosstalk ^3, 17, 19, 30-32^. Finally, relatively small changes in secreted factor levels, even if not statistically resolved at the protein expression level, may still be sufficient to drive measurable biological responses in sensitive downstream cell populations.

Overall, this work demonstrates that engineering design parameters—including geometry, control systems, and fluid mechanics—directly influence the biological fidelity of shear stress experiments. The presented bioreactor provides a versatile platform for investigating cardiovascular mechanobiology and establishes a framework for future studies integrating well-defined shear stimulation with co-culture systems and engineered tissues. Future work incorporating temporal profiling of mechanosensitive signaling and direct integration of co-culture within the device will further strengthen the quantitative linkage between shear stress inputs and downstream cellular responses.

## REFERENCES

1. Chistiakov, D.A., A.N. Orekhov, and Y.V. Bobryshev, Effects of shear stress on endothelial cells: go with the flow. Acta Physiol (Oxf), 2017. 219(2): p. 382–408, PMCID: PMID: 27246807.

2. Taber, L.A., J. Zhang, and R. Perucchio, Computational model for the transition from peristaltic to pulsatile flow in the embryonic heart tube. J Biomech Eng, 2007. 129(3): p. 441–9, PMCID: PMID: 17536912.

3. Yoshizumi, M., H. Kurihara, T. Sugiyama, F. Takaku, M. Yanagisawa, T. Masaki, and Y. Yazaki, Hemodynamic shear stress stimulates endothelin production by cultured endothelial cells. Biochem Biophys Res Commun, 1989. 161(2): p. 859–64, PMCID: PMID: 2660793.

4. Dewey, C.F., Jr., S.R. Bussolari, M.A. Gimbrone, Jr., and P.F. Davies, The dynamic response of vascular endothelial cells to fluid shear stress. J Biomech Eng, 1981. 103(3): p. 177–85, PMCID: PMID: 7278196.

5. Levesque, M.J. and R.M. Nerem, The elongation and orientation of cultured endothelial cells in response to shear stress. J Biomech Eng, 1985. 107(4): p. 341–7, PMCID: PMID: 4079361.

6. Uematsu, M., Y. Ohara, J.P. Navas, K. Nishida, T.J. Murphy, R.W. Alexander, R.M. Nerem, and D.G. Harrison, Regulation of endothelial cell nitric oxide synthase mRNA expression by shear stress. Am J Physiol, 1995. 269(6 Pt 1): p. C1371–8, PMCID: PMID: 8572165.

7. Walpola, P.L., A.I. Gotlieb, and B.L. Langille, Monocyte adhesion and changes in endothelial cell number, morphology, and F-actin distribution elicited by low shear stress in vivo. Am J Pathol, 1993. 142(5): p. 1392–400, PMCID: PMC1886930, PMID: 8494043.

8. Wang, N., H. Miao, Y.S. Li, P. Zhang, J.H. Haga, Y. Hu, A. Young, S. Yuan, P. Nguyen, C.C. Wu, and S. Chien, Shear stress regulation of Kruppel-like factor 2 expression is flow pattern-specific. Biochem Biophys Res Commun, 2006. 341(4): p. 1244–51, PMCID: PMID: 16466697.

9. Hove, J.R., R.W. Koster, A.S. Forouhar, G. Acevedo-Bolton, S.E. Fraser, and M. Gharib, Intracardiac fluid forces are an essential epigenetic factor for embryonic cardiogenesis. Nature, 2003. 421(6919): p. 172–7, PMCID: PMID: 12520305.

10. Jamison, R.A., C.R. Samarage, R.J. Bryson-Richardson, and A. Fouras, In vivo wall shear measurements within the developing zebrafish heart. PLoS One, 2013. 8(10): p. e75722, PMCID: PMC3790852, PMID: 24124507.

11. Stankovic, Z., B.D. Allen, J. Garcia, K.B. Jarvis, and M. Markl, 4D flow imaging with MRI. Cardiovasc Diagn Ther, 2014. 4(2): p. 173–92, PMCID: PMC3996243, PMID: 24834414.

12. Wong, A.K. L.L. P, N. Boroda, S.R. Rosenberg, and S.Y. Rabbany, A Parallel-Plate Flow Chamber for Mechanical Characterization of Endothelial Cells Exposed to Laminar Shear Stress. Cell Mol Bioeng, 2016. 9(1): p. 127–138, PMCID: PMC5629975, PMID: 28989541.

13. Franzoni, M., I. Cattaneo, B. Ene-Iordache, A. Oldani, P. Righettini, and A. Remuzzi, Design of a cone-and-plate device for controlled realistic shear stress stimulation on endothelial cell monolayers. Cytotechnology, 2016. 68(5): p. 1885–96, PMCID: PMC5023562, PMID: 26754843.

14. Arthurs, C.J., K.D. Lau, K.N. Asrress, S.R. Redwood, and C.A. Figueroa, A mathematical model of coronary blood flow control: simulation of patient-specific three-dimensional hemodynamics during exercise. Am J Physiol Heart Circ Physiol, 2016. 310(9): p. H1242–58, PMCID: PMC4867386, PMID: 26945076.

15. Madhavan, S. and E.M.C. Kemmerling, The effect of inlet and outlet boundary conditions in image-based CFD modeling of aortic flow. Biomed Eng Online, 2018. 17(1): p. 66, PMCID: PMC5975715, PMID: 29843730.

16. Saber, N.R., N.B. Wood, A.D. Gosman, R.D. Merrifield, G.Z. Yang, C.L. Charrier, P.D. Gatehouse, and D.N. Firmin, Progress towards patient-specific computational flow modeling of the left heart via combination of magnetic resonance imaging with computational fluid dynamics. Ann Biomed Eng, 2003. 31(1): p. 42–52, PMCID: PMID: 12572655.

17. Lemmens, K., V.F. Segers, M. Demolder, and G.W. De Keulenaer, Role of neuregulin-1/ErbB2 signaling in endothelium-cardiomyocyte cross-talk. J Biol Chem, 2006. 281(28): p. 19469–77, PMCID: PMID: 16698793.

18. Meyer, D., T. Yamaai, A. Garratt, E. Riethmacher-Sonnenberg, D. Kane, L.E. Theill, and C. Birchmeier, Isoform-specific expression and function of neuregulin. Development, 1997. 124(18): p. 3575–86, PMCID: PMID: 9342050.

19. Parodi, E.M. and B. Kuhn, Signalling between microvascular endothelium and cardiomyocytes through neuregulin. Cardiovasc Res, 2014. 102(2): p. 194–204, PMCID: PMC3989448, PMID: 24477642.

20. Peshkovsky, C., R. Totong, and D. Yelon, Dependence of cardiac trabeculation on neuregulin signaling and blood flow in zebrafish. Dev Dyn, 2011. 240(2): p. 446–56, PMCID: PMID: 21246662.

21. Polizzotti, B.D., B. Ganapathy, S. Walsh, S. Choudhury, N. Ammanamanchi, D.G. Bennett, C.G. dos Remedios, B.J. Haubner, J.M. Penninger, and B. Kuhn, Neuregulin stimulation of cardiomyocyte regeneration in mice and human myocardium reveals a therapeutic window. Sci Transl Med, 2015. 7(281): p. 281ra45, PMCID: PMC5360874, PMID: 25834111.

22. Cheng, S.W., E.S. Lam, G.S. Fung, P. Ho, A.C. Ting, and K.W. Chow, A computational fluid dynamic study of stent graft remodeling after endovascular repair of thoracic aortic dissections. J Vasc Surg, 2008. 48(2): p. 303–9; discusion 309–10, PMCID: PMID: 18644477.

23. Brooks, D.E., J.W. Goodwin, and G.V. Seaman, Interactions among erythrocytes under shear. J Appl Physiol, 1970. 28(2): p. 172–7, PMCID: PMID: 5413303.

24. Cherry, E.M. and J.K. Eaton, Shear thinning effects on blood flow in straight and curved tubes. Physics of Fluids, 2013. 25(7): p. 073104, PMCID: PMID:

25. Vennemann, P., K.T. Kiger, R. Lindken, B.C. Groenendijk, S. Stekelenburg-de Vos, T.L. ten Hagen, N.T. Ursem, R.E. Poelmann, J. Westerweel, and B.P. Hierck, In vivo micro particle image velocimetry measurements of blood-plasma in the embryonic avian heart. J Biomech, 2006. 39(7): p. 1191–200, PMCID: PMID: 15896796.

26. Cherry, E.M. and J.K. Eaton, Shear thinning effects on blood flow in straight and curved tubes. Physics of Fluids, 2013. 25(7), PMCID: PMID: WOS:000322521100026.

27. Bersell, K., S. Arab, B. Haring, and B. Kuhn, Neuregulin1/ErbB4 signaling induces cardiomyocyte proliferation and repair of heart injury. Cell, 2009. 138(2): p. 257–70, PMCID: PMID: 19632177.

28. Chatterjee, S. and A.B. Fisher, Mechanotransduction in the endothelium: role of membrane proteins and reactive oxygen species in sensing, transduction, and transmission of the signal with altered blood flow. Antioxid Redox Signal, 2014. 20(6): p. 899–913, PMCID: PMC3924805, PMID: 24328670.

29. Zheng, Q., Y. Zou, P. Teng, Z. Chen, Y. Wu, X. Dai, X. Li, Z. Hu, S. Wu, Y. Xu, W. Zou, H. Song, and L. Ma, Mechanosensitive Channel PIEZO1 Senses Shear Force to Induce KLF2/4 Expression via CaMKII/MEKK3/ERK5 Axis in Endothelial Cells. Cells, 2022. 11(14), PMCID: PMC9317998, PMID: 35883633.

30. Colliva, A., L. Braga, M. Giacca, and S. Zacchigna, Endothelial cell-cardiomyocyte crosstalk in heart development and disease. J Physiol, 2020. 598(14): p. 2923–2939, PMCID: PMC7496632, PMID: 30816576.

31. Kivela, R., K.A. Hemanthakumar, K. Vaparanta, M. Robciuc, Y. Izumiya, H. Kidoya, N. Takakura, X. Peng, D.B. Sawyer, K. Elenius, K. Walsh, and K. Alitalo, Endothelial Cells Regulate Physiological Cardiomyocyte Growth via VEGFR2-Mediated Paracrine Signaling. Circulation, 2019. 139(22): p. 2570–2584, PMCID: PMC6553980, PMID: 30922063.

32. Talman, V. and R. Kivela, Cardiomyocyte-Endothelial Cell Interactions in Cardiac Remodeling and Regeneration. Front Cardiovasc Med, 2018. 5: p. 101, PMCID: PMC6108380, PMID: 30175102.

